# SARS-CoV-2 Exploits Host Translation and Immune Evasion Pathways via Viral RNA–Host Protein Interactions

**DOI:** 10.1101/2025.03.07.642080

**Authors:** Kuerbannisha Amahong, Yuhong Liu, Zheng Zhang, Lin Tao, Aishe A. Sarshad, Feng Zhu

## Abstract

RNA viruses, including SARS-CoV-2, Influenza A Virus (IAV), Zika Virus, and Dengue Virus (DENV) pose serious global health challenges by manipulating host cellular mechanisms. SARS-COV-2, in particular exploits host translational machinery to enhance replication and evade immune response. Here, we investigated how SARS-CoV-2 circumvents host immune defenses through RNA - host protein interactions. By integrating multiple datasets, ClusterProfiler, KEGG, Reactome, WikiPathways, and Gene Ontology, we performed functional enrichment analyses on host protein interactions with SARS-CoV-2 RNA. Our results identified key pathways involved in viral replication, translation regulation, and immune evasion. Comparing SARS-CoV-2 interactomes from IAV, Zika, and DENV, we uncovered a subset of 275 common host proteins serving as promising targets for broad-spectrum antiviral strategies. Network analysis highlighted critical translation factors (EEF1A1, EIF4A1, EIF3H) and RNA-binding proteins (NCL, ILF3) as key nodes in viral replication. These findings provide insights into RNA virus pathogenesis and support the development of targeted therapeutics.

## INTRODUCTION

The emergence of RNA viruses such as Severe Acute Respiratory Syndrome Coronavirus 2 (SARS-CoV-2), Influenza A Virus (IAV), Zika Virus, and Dengue Virus (DENV) has posed serious global health challenges, widespread pandemics, significant loss of life and overwhelmed healthcare systems globally. The high variability and unpredictable interactions of RNA viruses with host cells complicate the development of vaccines and therapies ^1,2^. Therefore, understanding viral-host interactions is essential to elucidate viral life cycles and pathogenic mechanisms, to create effective therapeutic solutions ^3,4^.

The life cycle of RNA viruses involves complex biological processes including replication, transport, and the successful delivery of their RNA genomes into new host cells. These viruses lack the ability to encode all the proteins required for their replication, forcing them to rely on host cell molecules to compensate for the deficiencies. In response to viral invasion, host cells have evolved innate immune mechanisms that recognize viral RNAs and replication intermediates, triggering antiviral defenses ^5,6^.

The host system detects unconventional molecular signatures on viral RNA - such as 5’ triphosphates, improperly methylated cap structures, and double-stranded RNA (dsRNA) - to trigger an antiviral response within the cell, effectively suppressing viral gene expression ^7-9^. Among the first line sensors, Retinoic Acid-Inducible Gene I (RIG-I) and Melanoma Differentiation-Associated protein 5 (MDA5) receptors play key roles in detecting viral RNA ^10^. Additionally, endosomal Toll-Like Receptors 3 (TLR3) primarily detects dsRNAs ^11^. These receptors are highly expressed in immune cells, particularly dendritic cells, where their activation induces Toll-like receptor signaling, leading to the production of type I interferons and pro-inflammatory cytokines. This signaling cascade promotes dendritic cell maturation and initiates antiviral immunity ^12^. This rapid innate immune response establishes an antiviral state to restrict infection. On the other hand, RNA virus have also developed multiple strategies to evade host detection to endure viral replication and infectivity. For example, SARS-CoV-2 RNA hijacks host RNA binding proteins (RBP) such as LARPI, as an antiviral factor that normally would restrict viral RNA, effectively repurposing it to enhance viral replication ^13^. Similarly, the DENV virus interacts with the host splicing factor RBM10, deregulating host cell splicing and innate immune response to enhance viral replication ^14^.

In recent years, scientists have employed virus-RNA-protein interaction genomics to reveal how RNA viruses manipulate various host signaling pathways—such as inflammation, oxidative stress, apoptosis, and autophagy—to overcome immune defenses. By cross-referencing viral RNA interaction datasets with host protein databases, researchers have identified host proteins that are known drug targets and evaluated the effects of these compounds on viral infection. This strategy has revealed several cellular processes and signaling pathways, such as translation initiation and elongation, stress granule assembly, mTOR signaling, and interferon responses, whose drug inhibition can interfere with viral replication. For example, Schmidt et al. found that drug-induced inhibition of selected SARS-CoV-2 RNA interactome host proteins, including PPIA, ARP2/3, and ATP1A1, resulted in a dose-dependent reduction in viral replication ^15^. Kamel et al. observed that drug targeting of Heat Shock Protein 90 (HSP90) and Insulin-like Growth Factor 2 mRNA Binding Protein 1 (IGF2BP1) strongly inhibited SARS-CoV-2 protein synthesis, which was consistent with the antiviral effects of HSP90 ^16,17^. Furthermore, the interaction between IAV RNA and host protein DPP4 demonstrated that DPP4 is involved in activating several pathways, including triggering cytokine storms ^15^.

These interactions exhibit a dual role in both promoting and suppressing viral replication. For example, the interaction between SARS-CoV-2 RNA and host proteins ANGEL2 and La autoantigen, which are typically involved in mRNA stability and translation, is “hijacked” during viral infection to promote stability and efficient translation of viral mRNA ^16,17^. Moreover, some host RNA-binding proteins can recognize specific viral RNA sequences and activate immune responses, such as stimulating interferon signaling pathways to suppress viral replication and spread ^18^. Researches have also revealed how HCV and DENV manipulate inflammatory pathways, such as NF-κB, to promote their replication [12], and how DENV adjusts the redox state to interfere with oxidative stress responses ^19^, and how DENV adjusts the redox state to interfere with oxidative stress responses ^20^. Simultaneously, SARS-CoV-2 utilizes interactions with Bcl-2 family proteins to inhibit apoptosis ^21^, or uses autophagy to aid replication and assembly. These strategies not only enhance the viral spread but also help them evade immune surveillance. Overall, the inhibition of druggable targets identified from the SARS-CoV-2 RNA-protein interactome dataset confirms that these findings provide valuable resources for the development of antiviral therapies.

In this study we systematically explored the interactions between RNA genomes of SARS-CoV-2, IAV, Zika, and DENV, and host proteins using advanced enrichment analysis tools such as KEGG, Reactome, Wiki, and GO. By comparing host protein interaction information from multiple datasets, we found common host responses to these viral infections, as well as the unique roles of certain proteins in virus-host interactions. PPI network analysis further identified key host proteins interacting with these viral RNAs and examined their potential functions in the viral life cycle, particularly in viral replication and immune evasion strategies. These findings not only provide new insights into how hosts respond to RNA virus invasions but also lay a solid foundation for identifying new therapeutic targets and strategies against SARS-CoV-2. This research shows the value of multi-faceted analytical methods in elucidating the complex mechanisms by which host cells respond to viral invasion, providing scientific evidence for the development of future antiviral strategies (**Fig 1**).

**Figure 1.**
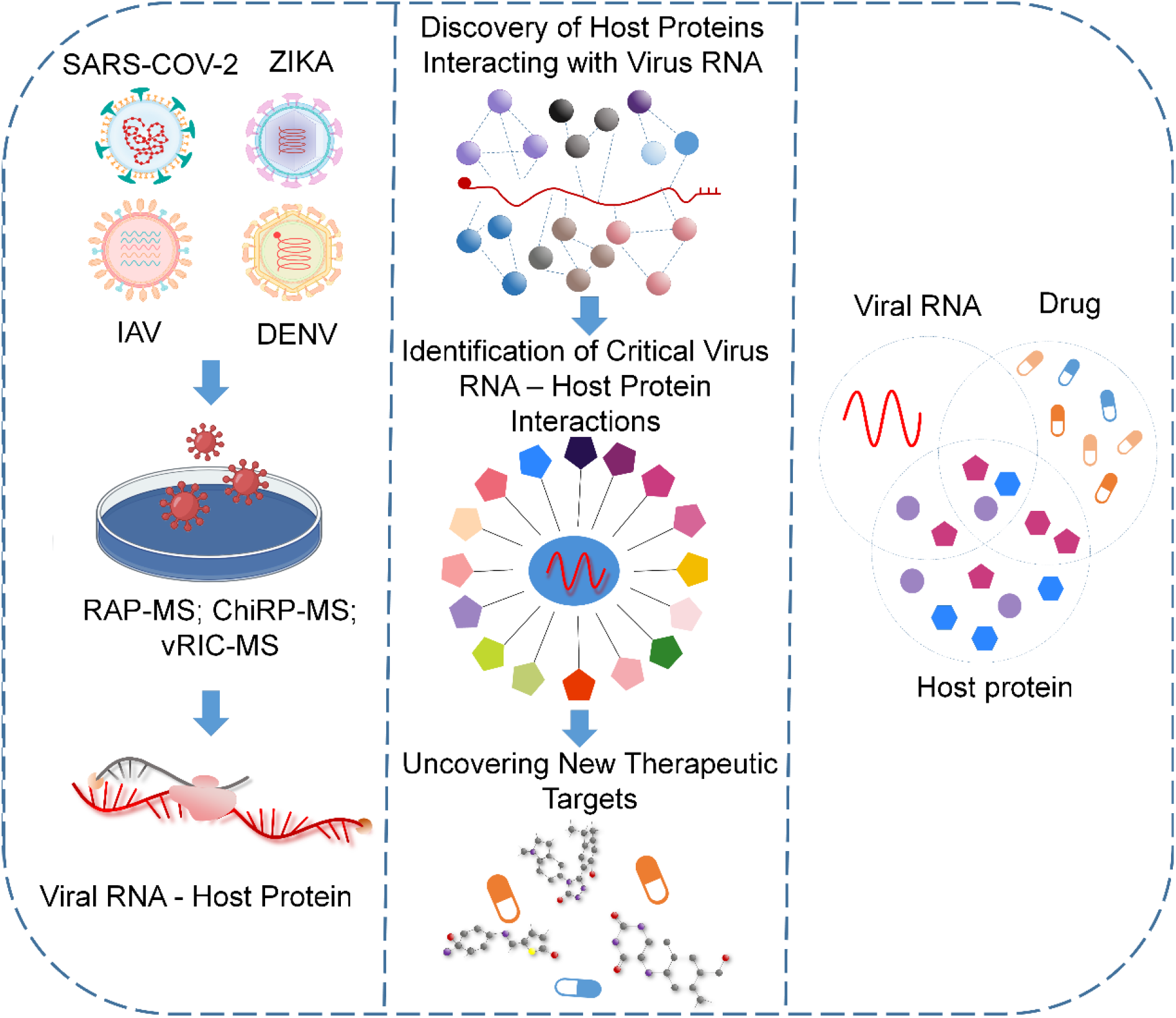
Overview of systematic analysis of viral RNA and host protein interaction data.

## Results and Discussions

### KEGG pathway analysis highlights RNA virus replication and immune evasion strategies through host protein interactions

The replication of SARS-CoV-2 and the production of progeny viruses rely on its ability to efficiently exploit host cell RNA stability, processing, localization, and translation regulatory mechanisms. Within the host cell, the successful recognition of the pathogen and the initiation of appropriate innate immune response pathways are critical to restraining the spread of viral infection ^22^. The top ranked KEGG pathway analysis revealed that these host proteins are associated with several key signaling pathways related to viral infection, including ‘Information processing in viruses,’ ‘Infectious disease: viral,’ ‘Immune system,’ ‘Translation,’ ‘Transcription,’ ‘Replication and repair,’ and ‘Folding, sorting, and degradation’ (**Fig 2A-H**). These findings highlight the complexity and diversity of host protein interactions with SARS-CoV-2 and also underscores the critical role of host proteins in the viral life cycle, particularly in the immune evasion mechanisms of the virus. Our analysis suggests that the functions of these host proteins extend far beyond supporting viral replication and dissemination; they are also implicated in the virus’s strategies to evade host immune responses. Therefore, a deeper understanding of the roles these host proteins play in the SARS-CoV-2 life cycle and immune evasion strategies is essential for uncovering the complexity of virus-host interactions and for identifying new therapeutic targets against SARS-CoV-2.

**Figure 2.**
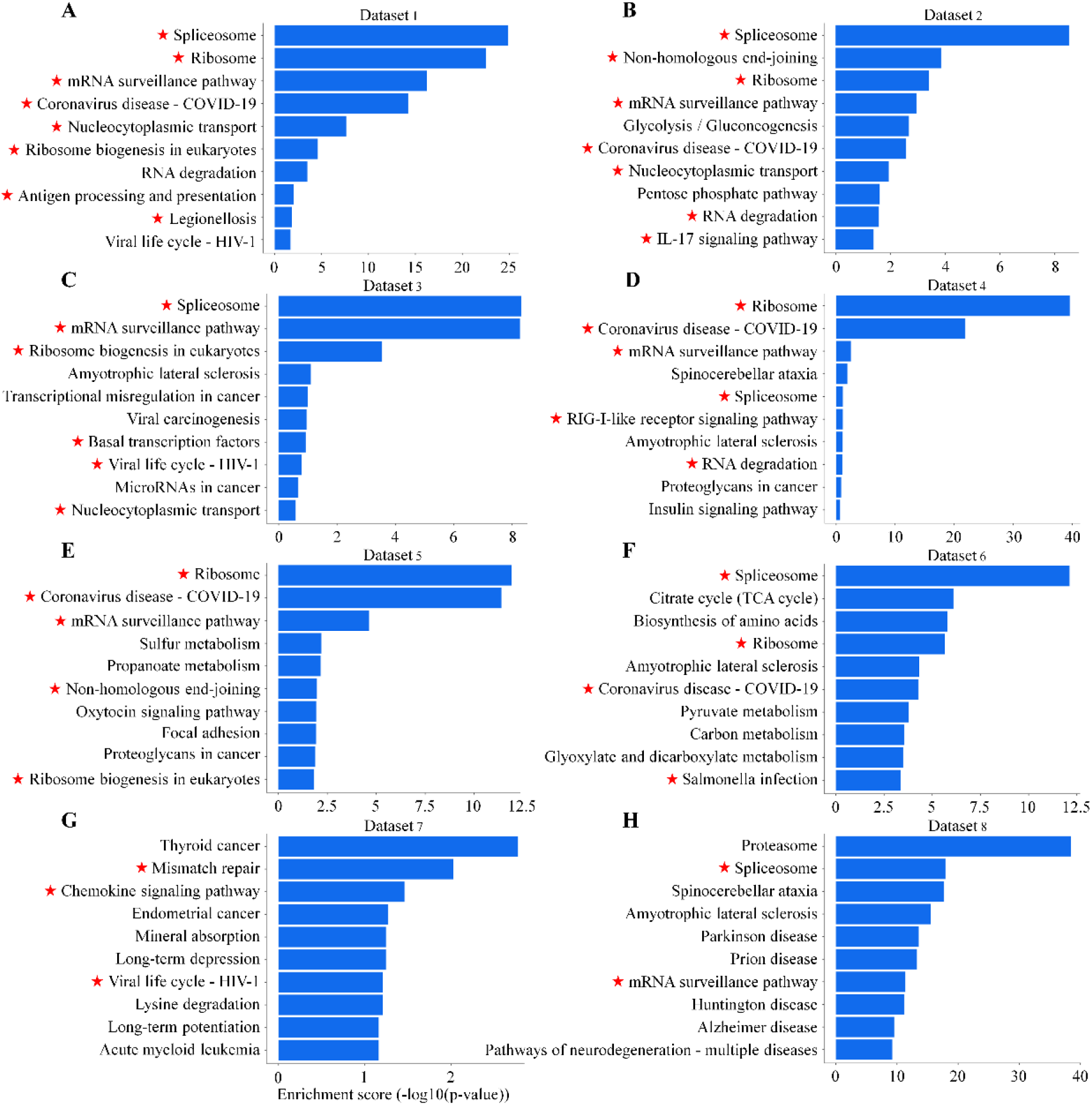
Enrichment analysis of KEGG pathways in eight datasets. (A–H) The bar graphs show top-ten enrichment pathways using host proteins that interacted with SARS-CoV-2 RNA genome in eight datasets according to enrichment score (−log10 (P-value)).

### SARS-CoV-2 hijacks and reprograms host translation machinery

SARS-CoV-2 employs a series of intricate strategies to interfere with and manipulate host protein functions ^23-25^. Here we used Reactome enrichment analysis to delve deeper into the biological functions of host proteins interacting with the SARS-CoV-2 RNA genome. Our analysis highlighted the importance of several key biological processes and immune-related signaling pathways (**Sup Fig 1A-H**). First, we found a significant enrichment in the translation process, particularly in the formation of the translation initiation complex and cap-dependent translation initiation. This finding underscores how SARS-CoV-2 utilizes host translation machinery to facilitate its protein synthesis. Moreover, the discovery of L13a-mediated translational silencing of ceruloplasmin expression demonstrates how the virus may influence host responses by modulating the translation of specific proteins. The analysis also highlighted the enrichment of the FGFR2 signaling pathway, indicating that SARS-CoV-2 may regulate host cell signaling through this pathway, thus promoting viral invasion and inflammatory responses. Notably, the enrichment of nonsense-mediated mRNA decay (NMD), particularly the NMD independent of the exon junction complex, suggests that SARS-CoV-2 may interfere with host mRNA surveillance mechanisms. Furthermore, the enrichment of cell cycle control and DNA damage response pathways, especially the p53-independent G1/S DNA damage checkpoint, suggests how viral infection could disrupt host cell cycle regulation and DNA repair mechanisms.

### SARS-CoV-2 exploits host RNA processing and metabolic pathways to evade immunity

Human innate immune defense mechanisms have evolved to detect and eliminate pathogens. Due to their relatively simple structures, viruses often evade early recognition by host cells. Over the course of evolution, viral nucleic acids have become one of the most powerful and reliable early indicators of infection. In the context of RNA virus detection, RNA sensing is a central mechanism in human innate immunity, and the efficacy of this sensing is critical for triggering an appropriate antiviral response. Despite the presence of various highly specialized receptors in human cells, which are designed to respond exclusively to pathogenic viral RNA, RNA viruses have evolved a range of strategies to avoid detection by the host’s interferon-mediated defense system. These viral evasion mechanisms are highly diverse, ranging from masking pathogenic RNA through terminal modifications to employing sophisticated techniques to deceive host cell RNA degradation enzymes, as well as hijacking basic metabolic pathways of the host cell ^26^. Our Wiki enrichment analysis (**Sup Fig 2A-H**) suggests that SARS-CoV-2 affects host mRNA processing, translation, and protein degradation, which may contribute to immune evasion and viral replication. The analysis showed that SARS-CoV-2 interferes with mRNA processing, translation factor activity, and the function of cytoplasmic ribosomal proteins, deeply affecting the host’s fundamental protein synthesis processes and preferentially promoting viral protein production ^27^. In our analysis, key translation factors and ribosome-related pathways, were enriched, indicating that the virus hijacks host translation machinery. This disruption may suppress host protein production while enhancing viral mRNA translation. Moreover, the virus manipulates specific cell signaling pathways, such as the VEGFA-VEGFR2 axis, to influence cell survival, proliferation, and angiogenesis, thereby enhancing its replication and spread while also interfering with the host’s inflammatory and immune responses ^28^.

Viral infection was also observed to induce changes in host cell metabolic pathways, which not only support the energy and biosynthetic precursors required for viral replication but may also lead to metabolic abnormalities associated with certain cancers ^29^. Our results also show enrichment in metabolic reprogramming in colon cancer, the TCA cycle, and cholesterol metabolism, suggesting that SARS-CoV-2 affects key metabolic processes. Although type I interferon signaling plays a crucial role in host antiviral responses ^30,31^, our analysis specifically highlights enrichment in the RIG-I-like receptor pathway, which is involved in viral RNA recognition, and host-cell autophagy, which may influence immune responses. These findings suggest that SARS-CoV-2 modulates metabolic and immune pathways to enhance its replication and persistence.

Recent studies have revealed a potential link between SARS-CoV-2 infection and neurodegenerative diseases such as Alzheimer’s Disease (AD) and Parkinson’s Disease (PD). Li et al. employed Mendelian randomization to explore the genetic correlation between COVID-19 and several neurodegenerative diseases, identifying a significant association between COVID-19 hospitalization and increased risk of Alzheimer’s disease ^32^. Another study focused on the shared molecular characteristics between coronavirus infections and neurodegenerative diseases, identifying common genes and pathways related to inflammation and stress responses present in both conditions. This study emphasized the potential of developing broad-spectrum drugs targeting these shared pathways, which could benefit both viral infections and neurodegenerative diseases ^33^. Consistently, our Wiki enrichment analysis also highlights enrichment in pathways related to Alzheimer’s disease and its associated mRNA effects, as well as the Parkin-ubiquitin proteasomal system pathway. These findings further support the idea that SARS-CoV-2 may influence neurodegenerative processes through disruptions in RNA regulation and protein degradation (**Sup Fig 2A-H**).

### GO analysis reveals SARS-CoV-2 exploitation of host mRNA processing, translation, and splicing machinery to facilitate viral replication

In this study, a thorough Gene Ontology (GO) analysis was conducted to explore how SARS-CoV-2 manipulates host cells at the levels of biological processes, cellular components, and molecular functions, revealing its complex strategies for replication and dissemination. At the level of biological processes, the enrichment of terms such as “Regulation of mRNA Metabolic Process” and “mRNA Processing” indicates that SARS-CoV-2 can interfere with the host’s mRNA metabolism and processing, which is crucial for the replication of viral RNA and the expression of host proteins (**Fig 3A**). By this mechanism, the virus preferentially translates its own proteins, thereby creating favorable conditions for its replication. The enrichment of “RNA Splicing” and “Regulation of RNA Splicing” suggests that the virus may influence the host’s splicing machinery, particularly through transesterification reactions, to affect the maturation of mRNA, subsequently regulating the expression of both host and viral proteins (**Fig 3A)**.

**Figure 3.**
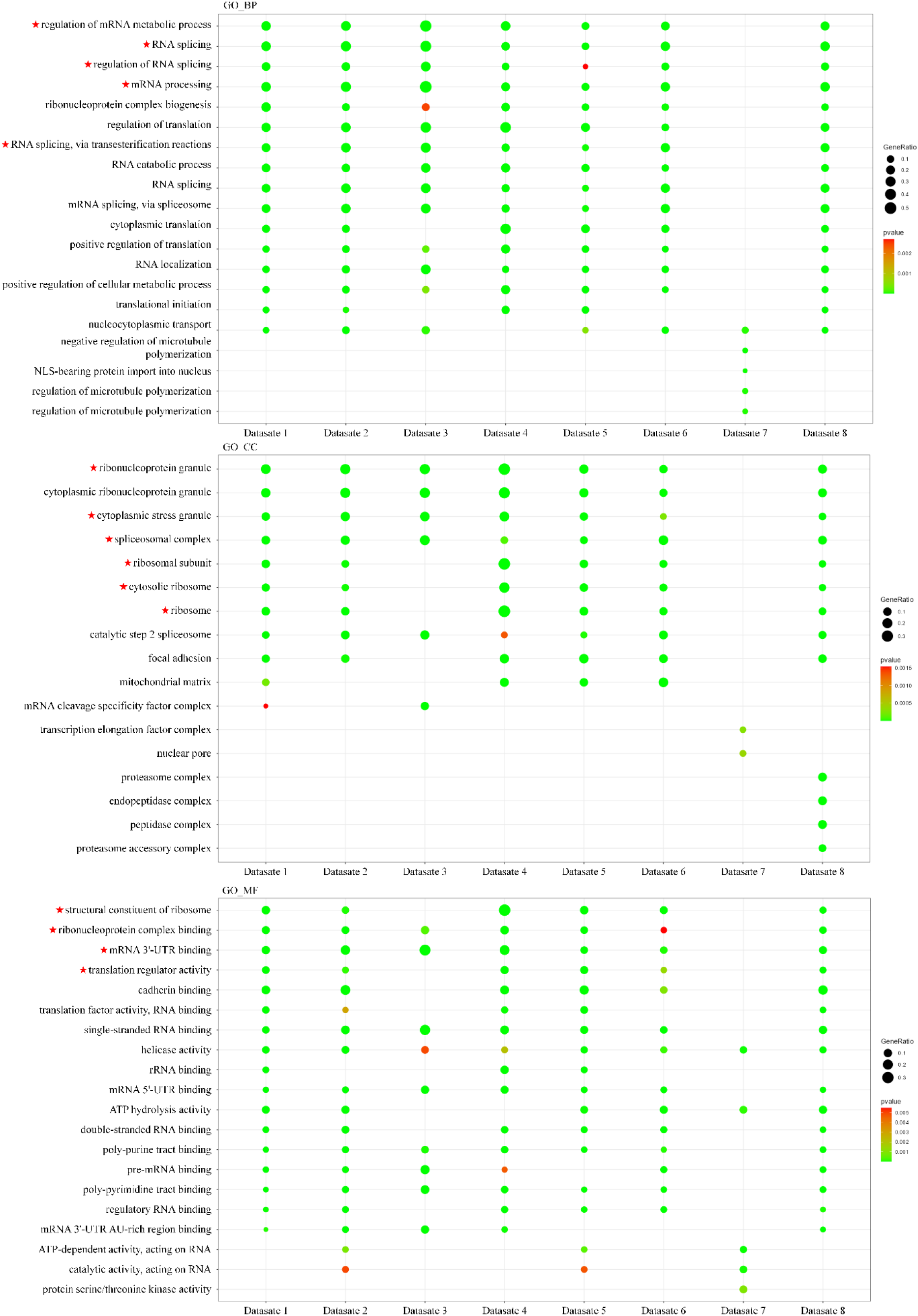
Gene Ontology (GO) analysis of host proteins interacting with the SARS-CoV-2 RNA genome across eight datasets. (A–C) Bar graphs depict the top ten most significantly enriched GO pathways for host proteins interacting with the SARS-CoV-2 RNA genome in each of the eight datasets, ranked by their enrichment scores (−log10 p-value).

In terms of cellular components, the enrichment of “Ribosomal Subunit” and “Ribosome” highlights the direct interaction between the virus and the host translation machinery, which could affect the synthesis of both viral and host proteins, particularly in structures such as the “Cytosolic Ribosome” and “Ribonucleoprotein Granule” (**Fig 3B**). Furthermore, the enrichment of “Spliceosomal Complex” and “Cytoplasmic Stress Granule” reflects how viral infection can affect RNA splicing and the host’s stress response to viral invasion. At the molecular function level, the emphasis on terms such as “Structural Constituent of Ribosome” and “Ribonucleoprotein Complex Binding” underscores the central role of these proteins in translation and their importance in the virus’s exploitation of host translation mechanisms. The enrichment of “mRNA 3’-UTR Binding” and “Translation Factor Activity, RNA Binding” further indicates that direct interactions between viral RNA and host proteins are crucial for the stability, translation, and intracellular localization of viral RNA (**Fig 3C**).

### Cross virus analysis reveals universal host responses and potential antiviral targets

This study aimed to elucidate the common response mechanisms of the host when confronted with infections from four RNA viruses: IAV, Zika, DENV, and SARS-CoV-2. In this regard, we first collected host protein interactome data with IAV, Zika, and DENV from the Rvvictor database, identifying 1545, 530, and 664 host proteins, respectively. These data were then compared with the host proteins interacting with SARS-CoV-2 (**Sup Table 1**). Through this integrated analysis, 275 key host proteins were identified to interact with all four viral RNAs, highlighting the universality of the host response and their potential roles in the viral infection process (**Fig 4**). To better understand the functional implication of these interactions, we performed functional annotation analysis using DAVID tool and the visualization through Cytoscape enhanced the understanding of these interactions (**Fig 4**). Notably, proteins identified as potential drug targets are highlighted in red drug symbols, demonstrating their function mechanisms. Additionally, we identified 35 potential drugs targeting 21 host proteins that interact with IAV, Zika, DENV, and SARS-CoV-2 RNA, providing valuable resources for antiviral therapeutic development.

**Figure 4.**
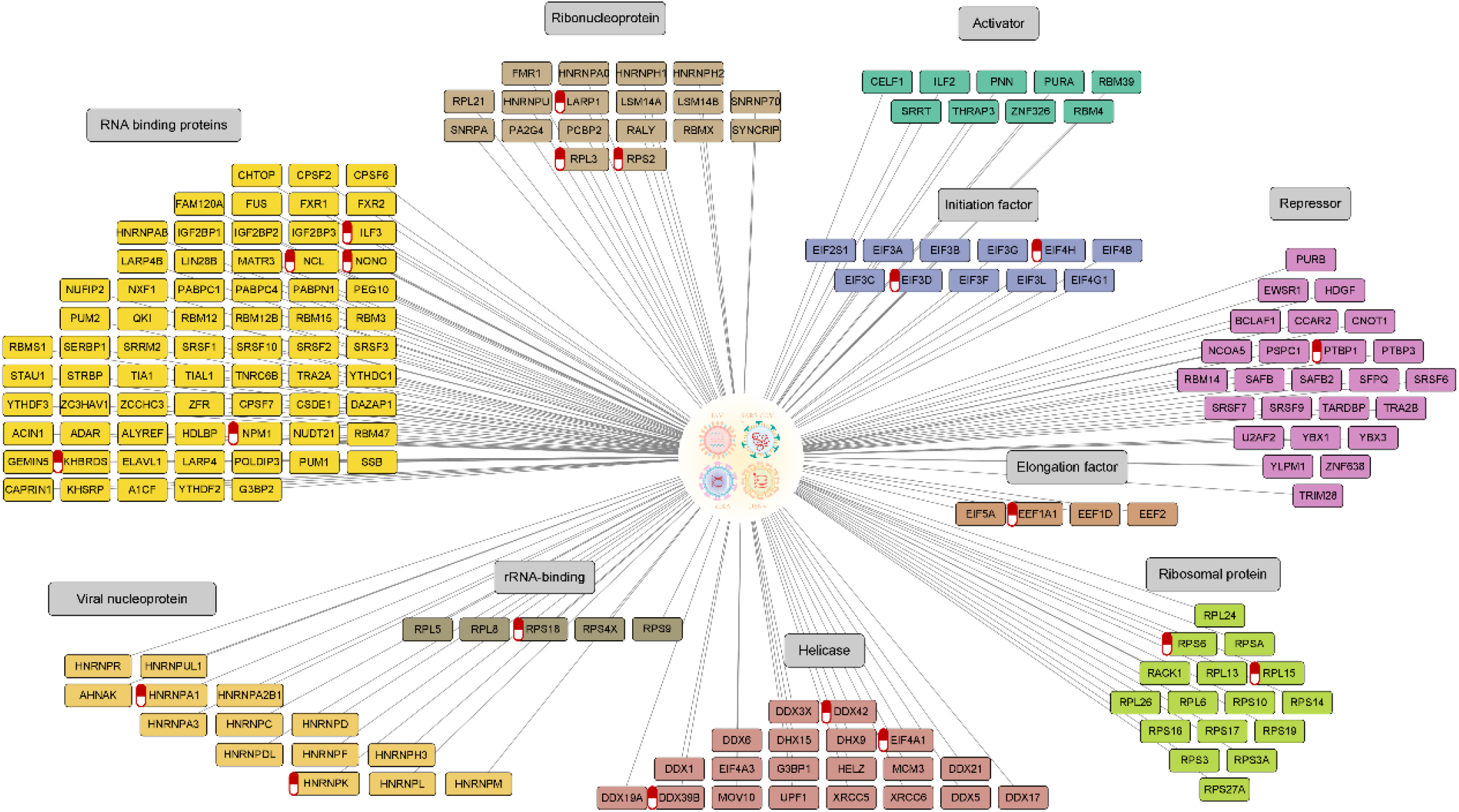
Comprehensive analysis and network visualization of host proteins interacting with IAV, Zika, DENV, and SARS-CoV-2

Collectively, these findings underscore the crucial role of host RNA binding proteins in RNA virus infections. The identified host proteins are involved in key cellular processes, including RNA-binding proteins, ribonucleoproteins, viral nucleoproteins, ribosomal proteins, helicases, initiation factors, repressor factors, activator factors, rRNA-binding proteins, and elongation factors (**Fig 4**). By deepening our understanding of these complex interactions, this study provides a theoretical foundation and new insights for the development of targeted antiviral strategies.

### Shared host protein networks across RNA viruses provide insights into broad-spectrum antiviral targets

A protein-protein interaction (PPI) analysis was conducted on the 275 shared host proteins interacting with four RNA viruses - IAV, Zika, DENV, and SARS-CoV-2 - using the STRING database. Interactions with a confidence score greater than 0.9 were identified (**Fig 5**). The results revealed critical roles of host proteins in the translation of viral RNA and the protein synthesis process, with particular emphasis on translation factors such as EEF1A1, EIF4A1, EIF4H, and EIF3H, as well as ribosomal proteins such as RPL3, RPS6, RPL8, RPL15, and RPS2. These proteins directly contribute to the efficient translation of viral mRNA, promoting viral replication. Notably, EEF1A1 has garnered significant attention for its importance in SARS-CoV-2 replication ^34^.

**Figure 5.**
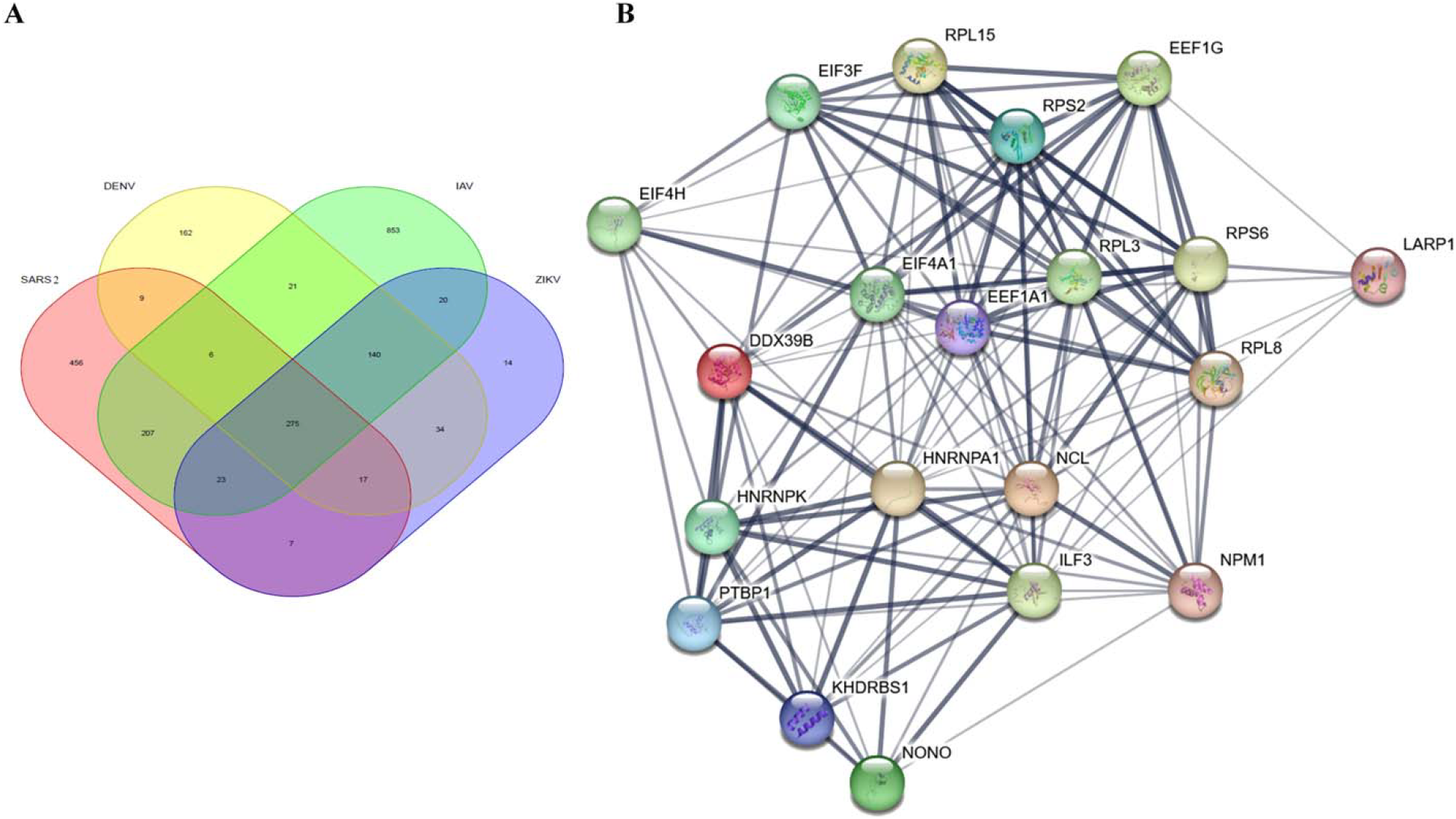
Conserved host protein interactions across four RNA viruses and PPI analysis. **A)** Venn diagram depicting the overlap of host proteins interacting with SARS-CoV-2, DENV, IAV, and ZIKV. The diagram illustrates the distribution of host proteins associated with each virus, highlighting 275 shared proteins that interact with all four RNA viruses. **B)** Protein-protein interaction (PPI) network analysis of 275 host proteins interacting with SARS-CoV-2, DENV, Influenza A Virus, and ZIKV through STRING Database. Key translation factors (EEF1A1, EIF4A1, EIF4H, EIF3H) and ribosomal proteins (RPL3, RPS6, RPL8, RPL15, RPS2) are highlighted due to their critical roles in viral mRNA translation. Proteins involved in RNA processing and stability (NCL, ILF3) and immune response regulation are also marked, revealing their potential roles in viral infection. The figure additionally mentions research progress on proteins as antiviral drug targets, such as EEF1A1 and Nucleolin, indicating these proteins as potential targets for future therapeutic strategies.

Additionally, proteins such as NCL, ILF3, NPM1, NONO, PTBP1, HNRNPK, and DDX39B play pivotal roles in RNA processing, including splicing, nuclear export, transport, and stability. This suggests that the virus may influence host RNA metabolism through these proteins, thereby disrupting normal cellular functions to facilitate viral replication ^35^. In the context of immune response and regulation, proteins such as ILF3 and NCL, which participate in the interferon signaling pathway, are critical in combating viral infections ^36^. The recurrent presence of these host proteins in infections caused by different RNA viruses highlights the universal dependency of host cell mechanisms, offering new perspectives for identifying broad-spectrum antiviral targets. Specifically, EEF1A1, Nucleolin, EIF4A1, and ILF3 play key roles in viral replication and serve as potential targets for antiviral drugs. Small-molecule inhibitors targeting EEF1A1 may serve as effective tools to block viral protein translation ^37^. Therapeutic strategies targeting Nucleolin have shown promise in blocking viral interactions ^38^, while the regulation of EIF4A1 and ILF3 is considered a potential antiviral treatment target ^39,40^. These findings provide a theoretical foundation and new insights into the understanding of how viruses exploit host cell mechanisms for replication and offer valuable directions for the development of targeted antiviral strategies.

## Conclusion

In this study we provide an in-depth analysis of the interactions between SARS-CoV-2 and host proteins, exploring viral replication, immune evasion, and their effects on related signaling pathways, thereby offering crucial insights for the development of effective antiviral therapeutic strategies. Functional enrichment analyses, including KEGG, Reactome, Wiki, and GO, conducted using tools such as Cluster Profiler, have identified key pathways related to the viral life cycle and immune evasion, highlighting the central role of host proteins in these processes. Our findings suggest that SARS-CoV-2 effectively promotes its replication and immune evasion through interactions with host factors involved in RNA stability, processing, localization, and translation regulation. Pathways identified in the KEGG analysis, such as “Information Processing in Viruses,” “Viral Diseases,” and “Immune System,” emphasize the pivotal role of host proteins in the virus’s immune evasion mechanisms.

Through the cross-virus host protein interaction network analysis, this study further revealed the common response mechanisms of the host cell when confronted with infections from RNA viruses such as SARS-CoV-2, IAV, Zika, and DENV. The 275 shared host proteins that interact with the RNA of all four viruses demonstrate the universal dependency of host cell mechanisms, providing new avenues for identifying broad-spectrum antiviral targets. Furthermore, the functional classification and visualization of interaction networks highlight the diversity and complexity of host protein responses during RNA virus infections. Notably, the study of host proteins such as EEF1A1, Nucleolin, EIF4A1, and ILF3 confirms that these datasets offer valuable resources for the development of antiviral therapies.

In summary, we provide an understanding of how SARS-CoV-2 exploits host cell mechanisms to facilitate replication and evade immune surveillance, offering a solid scientific foundation for the development of new antiviral therapeutic strategies. Future studies should further explore the detailed interaction mechanisms between host proteins and SARS-CoV-2 to aid in the development of targeted treatments for SARS-CoV-2. This will provide more effective therapeutic options for combating COVID-19 and other RNA virus infections.

## Materials and Methods

### Extraction of host proteins interacting with the SARS-CoV-2 RNA

Eight datasets from the CovInter database were selected ^41^, encompassing SARS-CoV-2 strains from various regions worldwide, including the UK (hCoV-19/England/02/2020), France (hCoV-19/France/IDF-220-95/2020), South Korea (hCoV-19/South Korea/KCDC03/2020), Germany (virus strains isolated from the University of Regensburg Clinical Microbiology and Hygiene Department), China (hCoV-19/IPBCAMS-YL01/2020, hCoV-19/Wuhan-Hu-1/2019), and USA (hCoV-19/USA/WA1/2020). The dataset from these studies employed a range of experimental techniques, such as RNA antisense purification coupled with mass spectrometry (RAP-MS), Label-Free Quantification (LFQ), Liquid Chromatography with Tandem Mass Spectrometry (LC-MS/MS), Tandem Mass Tag (TMT) labeling, RNA purification and chromatin isolation techniques combined with mass spectrometry (ChIRP-M/S), and RNA-protein interaction detection assays (RaPID), and successfully identifying 65 to 332 human proteins that potentially interact with SARS-CoV-2 RNA (**Table 1**).

**Table 1.**
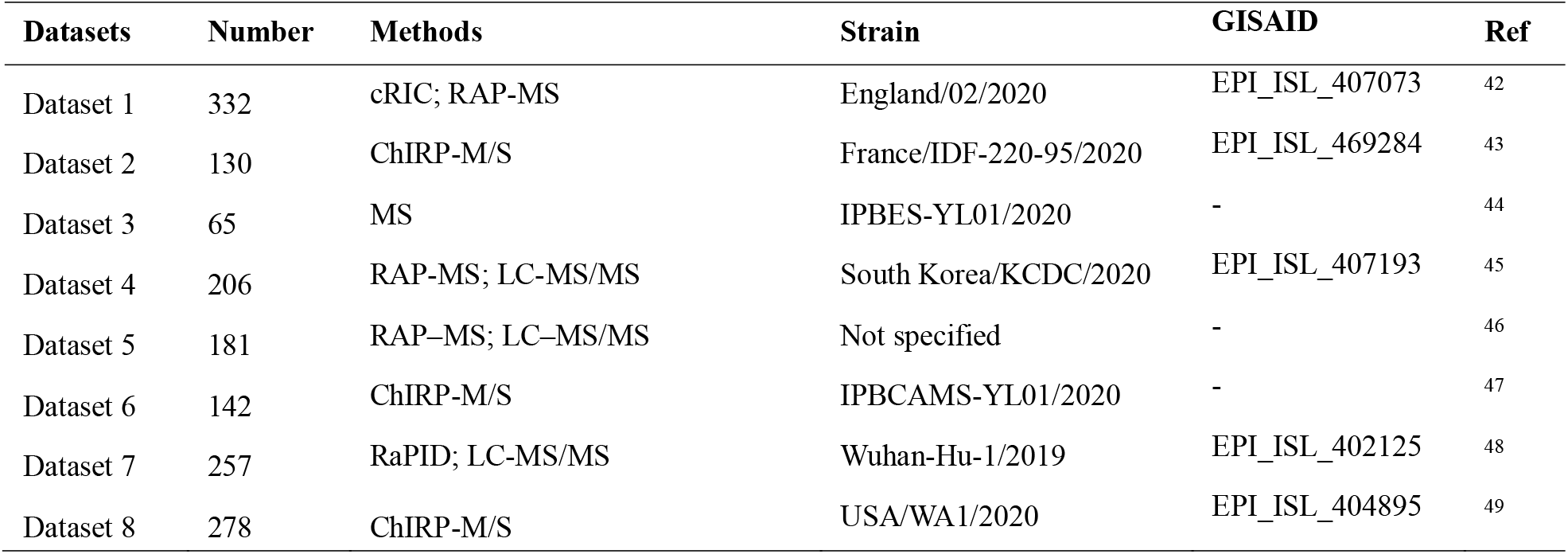
Eight datasets on host protein interactions with the SARS-CoV-2 RNA.

| Datasets | Number | Methods | Strain | GISAID | Ref |
| --- | --- | --- | --- | --- | --- |
| Dataset 1 | 332 | cRIC; RAP-MS | England/02/2020 | EPI_ISL_407073 | 42 |
| Dataset 2 | 130 | ChIRP-M/S | France/IDF-220-95/2020 | EPI_ISL_469284 | 43 |
| Dataset 3 | 65 | MS | IPBES-YL01/2020 | - | 44 |
| Dataset 4 | 206 | RAP-MS; LC-MS/MS | South Korea/KCDC/2020 | EPI_ISL_407193 | 45 |
| Dataset 5 | 181 | RAP-MS; LC-MS/MS | Not specified | - | 46 |
| Dataset 6 | 142 | ChIRP-M/S | IPBCAMS-YL01/2020 | - | 47 |
| Dataset 7 | 257 | RaPID; LC-MS/MS | Wuhan-Hu-1/2019 | EPI_ISL_402125 | 48 |
| Dataset 8 | 278 | ChIRP-M/S | USA/WA1/2020 | EPI_ISL_404895 | 49 |

### Functional enrichment analysis of host proteins interacting with SARS-CoV-2 RNA

Functional enrichment analysis was performed on the eight datasets using the ClusterProfiler software (version 4.4.4) ^1^. This analysis focused on three key areas: KEGG pathways, the Reactome database, and Gene Ontology (GO). The results were generated using the plotting package in the R environment (version 4.2.1).

### Comprehensive identification and network analysis of cross-species RNA virus-host protein interactions

To explore the common interaction mechanisms between IAV, Zika, DENV, and SARS-CoV-2 viruses and host proteins, host protein interaction data for IAV, Zika, and DENV were collected from the Rvvictor database. A total of 1545, 530, and 664 host proteins interacting with these three viruses were identified, respectively. By comparing these with the host proteins interacting with SARS-CoV-2, key host proteins common to all four viruses were selected. Subsequently, functional annotation analysis was performed using the DAVID tool, and the interaction network was visualized with Cytoscape software.

### Drug repurposing strategy for host protein targets in cross-virus interactions

To identify potential drugs targeting host proteins involved in the interactions with multiple viruses, a literature review was conducted, using keyword combinations such as “host protein name + drug” and “SARS-CoV-2 + drug repurposing” to search for relevant studies. This search successfully identified 35 drugs targeting 21 host proteins involved in interactions with IAV, Zika, DENV, and SARS-CoV-2 RNA.

### Protein-protein interaction analysis

Protein-protein interaction (PPI) analysis was conducted using the STRING database (https://string-db.org/) to investigate interactions among 275 shared host proteins, to identify critical interaction points in the viral infection and replication process. A confidence threshold of >0.9 was selected to ensure the high accuracy and reliability of the results, thus accurately pinpointing interactions that have a decisive impact on the viral life cycle.

## Data availability

The datasets used in this study is provided in Supplementary Table1.

## Acknowledgments

We thank the authors who generated the datasets used in this study.

## Author contributions

K.A; conceptualization, data curation, formal analysis, investigation, methodology, visualization, writing – original draft. Y.L; data curation, investigation. Z.Z; data curation, formal analysis, investigation. L.T; conceptualization, methodology. A.A.S; and F.Z; project administration, supervision. All authors contributed to the manuscript

## Competing Interests

The authors declare no competing interests.

## Notes

### Competing Interest Statement

The authors have declared no competing interest.

