## Supplementary material for "SARS-CoV-2 Exploits Host Translation and Immune Evasion Pathways via Viral RNA–Host Protein Interactions": x

**Supplemental Figures**

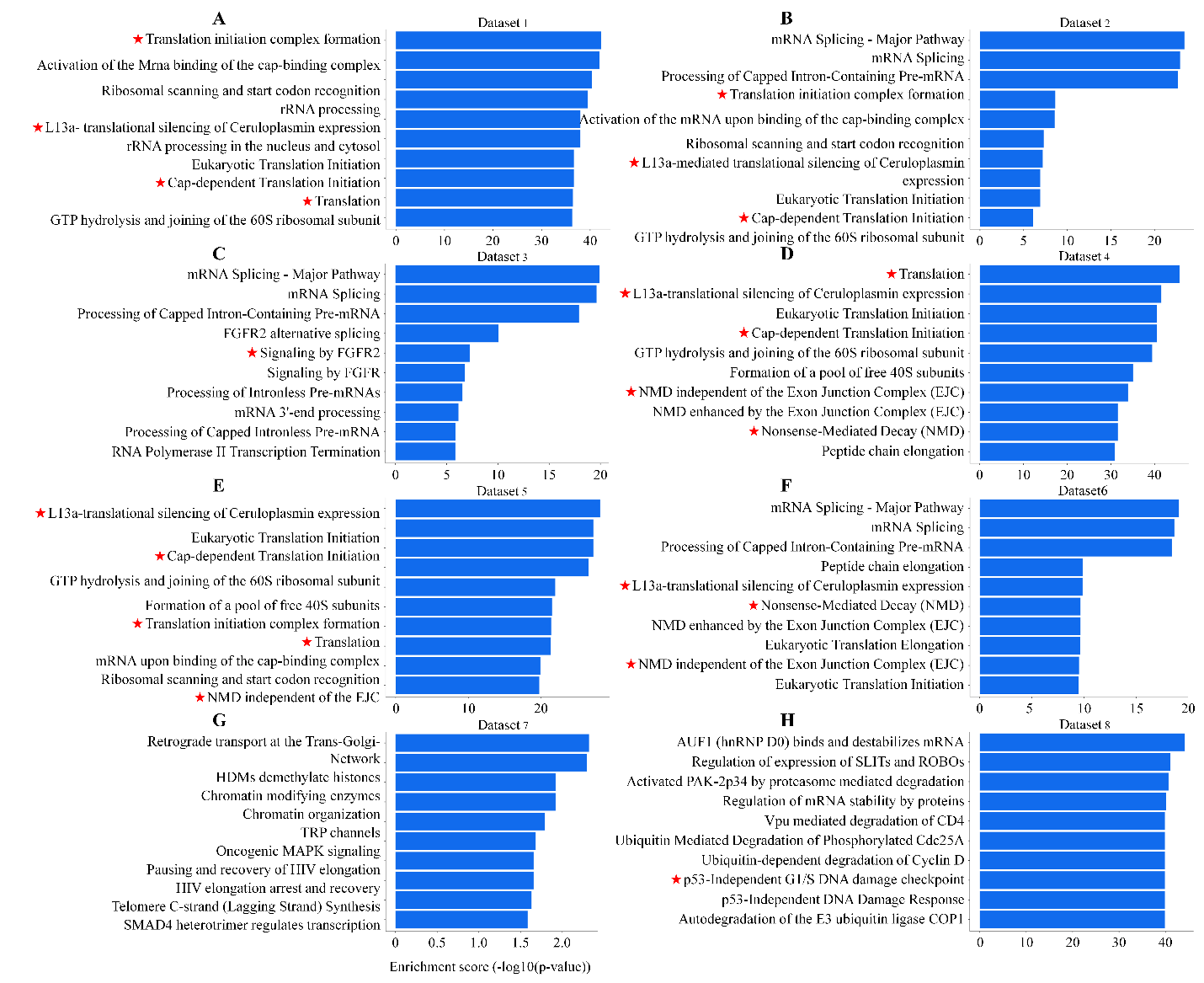

**Supplemental Figure 1.** Enrichment analysis of Reactome pathways in eight datasets. (A–H) Bar graphs show top-ten pathways of enrichment using host proteins that interacted with SARS-CoV-2 RNA genome in eight datasets according to Enrichment score (−log10 (P-value)).

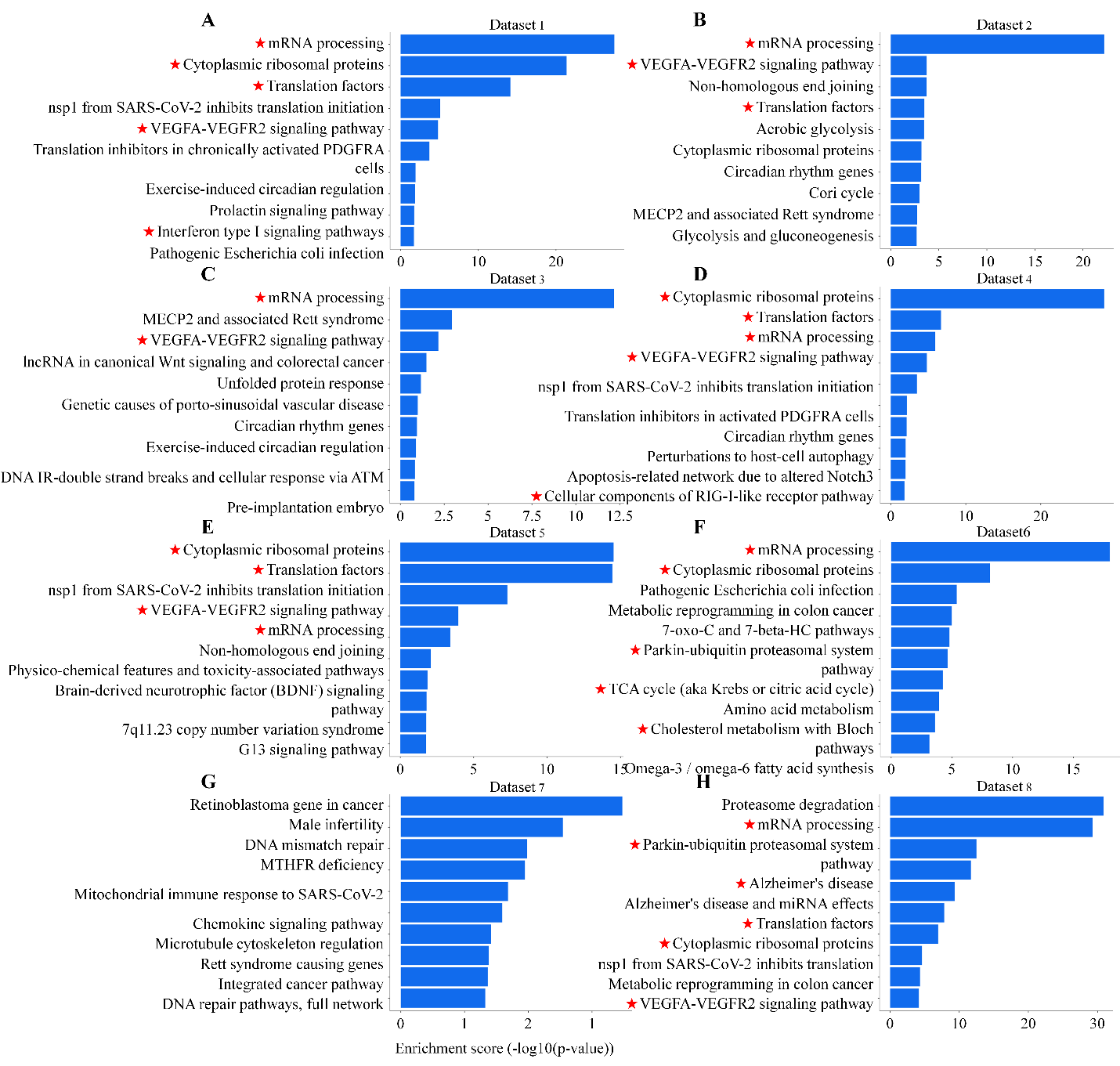

**Supplemental Figure 2** Enrichment analysis of Wiki pathways in eight datasets. (A–H) Bar graphs show top-ten pathways of enrichment using host proteins that interacted with SARS-CoV-2 RNA genome in eight datasets according to Enrichment score (−log10 (P-value)).
